# Cytochrome expression shifts in *Geobacter sulfurreducens* to maximize energy conservation in response to changes in redox conditions

**DOI:** 10.1101/2022.05.22.492868

**Authors:** Ethan Howley, Rosa Krajmalnik-Brown, César I. Torres

**Affiliations:** Biodesign Swette Center for Environmental Biotechnology, Arizona State University – Tempe, AZ; School of Sustainable Engineering and the Built Environment, Arizona State University – Tempe, AZ; Biodesign Center for Health Through Microbiomes, Arizona State University – Tempe, AZ; School for Engineering of Matter, Transport, and Energy, Arizona State University – Tempe, AZ

## Abstract

Previous studies have identified that *Geobacter sulfurreducens* has three different electron transfer pathways for respiration, and it switches between these pathways to adapt to the redox potential of its electron acceptor. However, only a small fraction of the electron carriers from each pathway have been identified. In this study, we combined electrochemical and gene expression data to identify electron carriers associated with each of the three pathways in the inner membrane, periplasm, outer membrane, and exterior of the cell. We demonstrate that it is not just the electron acceptor redox potential that controls pathway expression in *G. sulfurreducens*. Our method combining electrochemical modeling and transcriptomics could be adapted to better understand electron transport in other electroactive organisms with complex metabolisms.

**Graphical Abstract:** 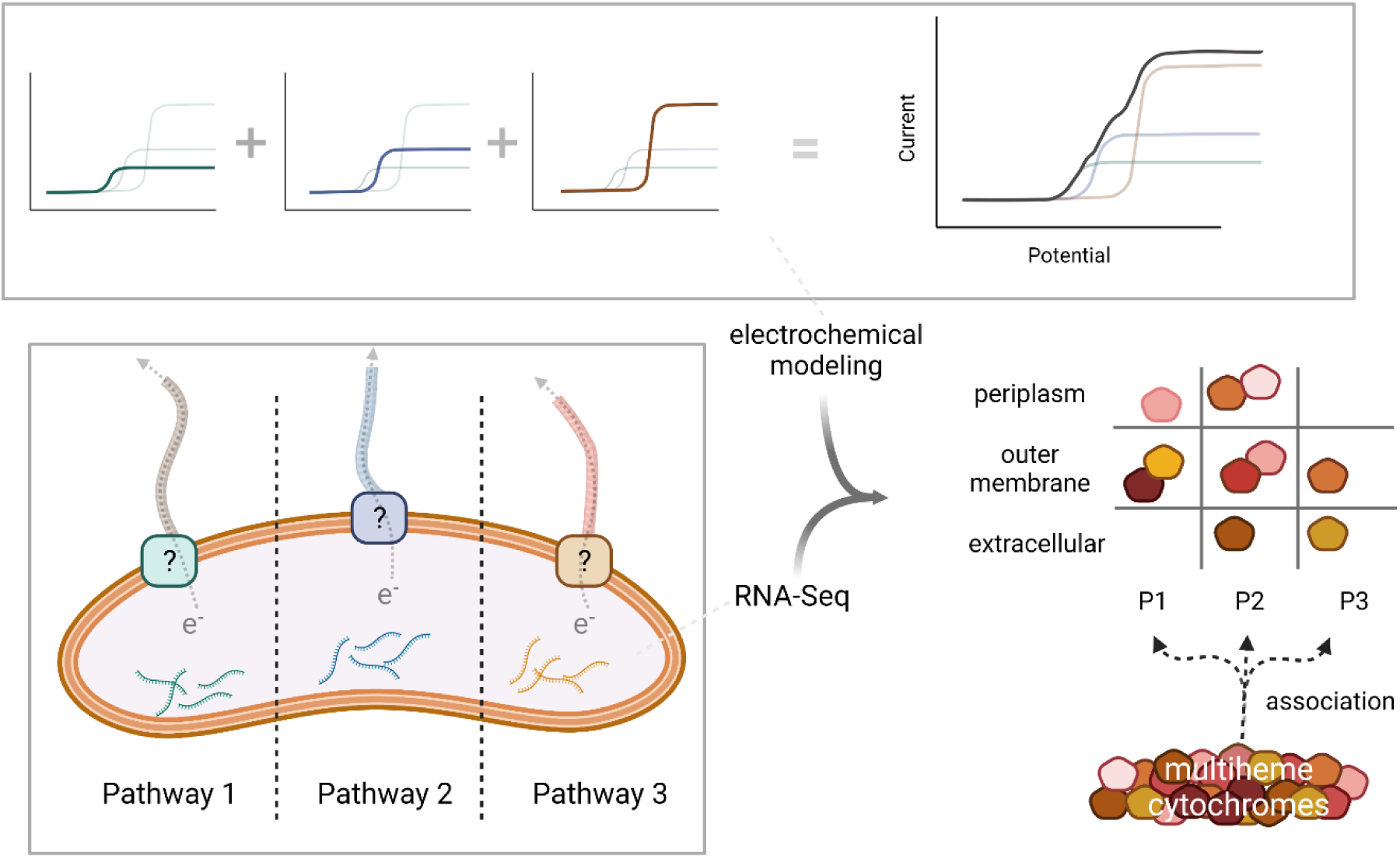

Graphical abstract was created with Biorender.com

## Introduction

*Geobacter sulfurreducens* is an anaerobic, gram negative, dissimilatory metal reducing bacterium (DMRB) that is important in natural systems as an iron reducer as well as in engineered microbial electrochemical systems as an anode-respiring bacterium (ARB) (Bond and Lovley 2003; Caccavo et al. 1994). It can efficiently reduce various metal oxides with a wide range of redox potentials (Levar et al. 2017; Pat-Espadas et al. 2013; Shelobolina et al. 2007), and this flexibility extends to respiring to an anode where *G. sulfurreducens* is capable of adapting to a range of poised anode potentials (Gao et al. 2018; Yoho, Popat, and Torres 2014).

The genome of *G. sulfurreducens* contains genes for over 100 putative c-type cytochromes, with more than 70 of those containing multiple heme-binding motifs (Ding et al. 2008). These multiheme cytochromes have been the target of many studies into electron transfer in *G. sulfurreducens*, but only a few have been definitively linked to a precisely explained function (Salgueiro et al. 2022). Even well-characterized cytochromes, such as the periplasmic cytochrome PpcA (Pessanha et al. 2006) or the outer membrane complex OmcB (Liu et al. 2015), are often known to participate in certain respiratory conditions, but their specific electron donor/acceptor are unknown. As such, the specific electron transport chain pathway for *G. sulfurreducens* has been extensively speculated but not well elucidated (Bonanni, Massazza, and Busalmen 2013; Santos et al. 2015).

Early transcriptomic and proteomic studies on *G. sulfurreducens* focused on comparing growth with different electron acceptors, particularly between soluble (e.g., fumarate, iron citrate) and solid ones (e.g. anodes, iron oxides). Through these studies, several cytochromes have been confirmed to be important in extracellular respiration. For example, OmcB, OmcS, OmcZ, and other cytochromes were implicated in extracellular electron transfer because their gene transcripts were more abundant in conditions using a solid terminal electron acceptor relative to an insoluble one (Holmes et al. 2006; Nevin et al. 2009). It has been hypothesized that long-range electron transfer in *G. sulfurreducens* was due to some combination of pili and associated cytochromes like OmcZ (Malvankar et al. 2011; Nevin et al. 2009). Recent cryo-electron microscopy studies have shown that the nanowires that *G. sulfurreducens* produce are actually composed of repeating subunits of OmcS or OmcZ, and differences in strain and electron acceptor can alter which protein is dominantly expressed (Wang et al. 2019; Yalcin et al. 2020).

More recently, researchers have discovered three elements at the inner membrane that are required for *G. sulfurreducens* to respire at certain potentials, independent of the electron acceptor used. Deleting the inner membrane cytochrome CbcL prevents *G. sulfurreducens* from respiring at electrode potentials below -0.1 V vs. SHE, and deleting the inner membrane cytochrome ImcH prevents *G. sulfurreducens* from respiring at potential above -0.1 V vs. SHE (Levar et al. 2017; Levar, Chan, and Mehta-kolte 2014; Zacharoff, Chan, and Bond 2016).

Recently, the protein complex CbcAB has been shown to facilitate respiration at potentials below -0.21 V vs. SHE (Joshi, Chan, and Bond 2021). At the moment, it is unknown how pathway switching occurs, and what other proteins are involved in each pathway. Nonetheless, these studies have shifted the perspective on how pathways are used in *G. sulfurreducens*, where instead of the type of electron acceptor, the potential of such acceptor plays a major role in pathway utilization. By using an anode as the electron acceptor, we can closely control potential and directly measure respiration in real time.

Previous work has also investigated how *G. sulfurreducens* adapts to electron acceptors with different potentials through the use of electrochemical signals (Peng and Zhang 2017; Richter et al. 2009; Zhu, Yates, and Logan 2012). Our group used electrochemical techniques to identify and characterize two distinct electrochemical responses dependent on anode potential and hypothesized that *G. sulfurreducens* is using different electron transfer pathways to efficiently adapt to different potentials (Yoho, Popat, and Torres 2014). These distinct signals were later associated to the pathways in which ImcH and CbcL are present (Levar et al. 2014, 2017). The relative magnitude of the signals associated to these pathways would shift within an hour of a potential shift, suggesting a dynamic and complex optimization of respiratory pathways by *G. sulfurreducens*. Anode-respiring biofilms of *G. sulfurreducens* exhibit different electrochemical responses during cyclic voltammetry based on the anode potential they have been acclimated to, providing an additional tool to differentiate between biofilms grown at different potentials.

In order for *G. sulfurreducens* to efficiently reduce solid electron acceptors at a wide range of redox potentials while conserving energy, it must have a mechanism to change its electron transport chain. Each cytochrome has its own distinct redox potential due to heme orientation and ligands within the overall protein (Pessanha et al. 2006; Pokkuluri et al. 2011), and therefore has an optimal potential (or range of potentials) under which it will accept and donate electrons in a pathway. If the terminal electron acceptor changes potential significantly, the terminal electron transfer protein may no longer be able to donate electrons and a different terminal electron acceptor will be required to continue respiration. Thus, we hypothesized that a shift in the potential of the electron acceptor would not only require shifts in inner-membrane cytochromes, but throughout the whole respiratory pathway. We designed this study to attempt to find periplasmic, outer membrane, and/or extracellular proteins that are associated with *G. sulfurreducens* respiration at different redox potentials.

We studied wild type *Geobacter sulfurreducens PCA* grown on anodes using electrochemistry and mRNA expression to identify genes that are likely involved in different electron transfer pathways. With cyclic voltammetry (CV) on *G. sulfurreducens* biofilms, we can identify if different pathway signals are present in biofilms grown at different anode potentials, and what typical anode potentials each pathway is associated with. We associated these CV signals with mRNA-Seq transcriptomics to correlate gene expression with the electrochemical data and find genes that may be associated with adaptation to different electron acceptor potentials. Since previous studies have identified the inner membrane elements associated with each pathway, we closely examined the periplasmic, outer membrane, and extracellular electron-transfer proteins that might shift expression as a function of anode potential.

## Results and Discussion

When studying ARB, we can directly measure respiration via chronoamperometry while varying anode potential to create different respiratory conditions. Anode biofilm cyclic voltammograms (CV) on biofilms of *G. sulfurreducens* can be modeled with the Nernst-Monod expression (Torres et al. 2008, 2010) which assumes a single enzymatic step is rate limiting. Additionally, in cases where multiple pathways are being utilized, multiple Nernst-Monod functions have been used (Yoho, Popat, and Torres 2014).

Knockout studies have shown that *G. sulfurreducens* has at least three different electron transfer pathways that are each active in only a certain electron acceptor range, and the switch between pathways is likely facilitated by inner membrane cytochromes (Levar et al. 2017). The pathway containing the inner membrane cytochrome ImcH is used at potentials higher than -0.1 V vs.

SHE, while the pathway containing the cytochrome CbcL is active at potentials lower than -0.1 V vs. SHE (Levar et al. 2014, 2017). Recently, a third pathway was discovered that contains the *bc*-type cytochrome CbcBA which is required for respiration at potentials below -0.21 V vs. SHE (Joshi, Chan, and Bond 2021). We chose the anode potentials in this experiment (−0.17 V, -0.07 V, -0.01 V vs. SHE) in an attempt to create conditions that would require different known pathways in *G. sulfurreducens*, and our modeling approach assumes three distinct electrochemical signals in attempt to represent these pathways.

### Multiple pathway signals fit to a Nernst-Monod model

We were able to approximately fit CVs of the anode biofilm conditions used in this study by adding Nernst-Monod signals with three midpoint potentials: −0.10, −0.15, and −0.227 V vs. SHE; which were selected based on preliminary nonlinear fitting where midpoint was optimized in different conditions (Figure 1). We call these signals High for −0.10 V vs. SHE, Medium for −0.15 V vs. SHE, and Low for −0.227 V vs. SHE. The additive approach to the fitting assumes the signals are utilized simultaneously within the biofilm and cells are not changing expression significantly during the scan. This assumption should be valid with the fast-scan CVs (5 mV/s) we used for fitting, while slower scans are shown to result in pathway shifts within the scan itself (Yoho, Popat, and Torres 2014). With nonlinear fitting, we minimized the error in each fitting by varying the j_max_ coefficient for each E_KA_. In some cases, such as the Acetate −0.17 V growth condition, fitting had a higher error due to a peak in current density that cannot be estimated by the steady-state assumption of the Nernst-Monod (Torres et al. 2008, 2010). We averaged the fractional contribution of each of the coefficients from biological replicates of the same condition to estimate the contribution of each pathway in that condition (Table 1). Fittings were tested in CVs and the derivative of the CV, having similar outcomes for the fitting.

**Figure 1:**
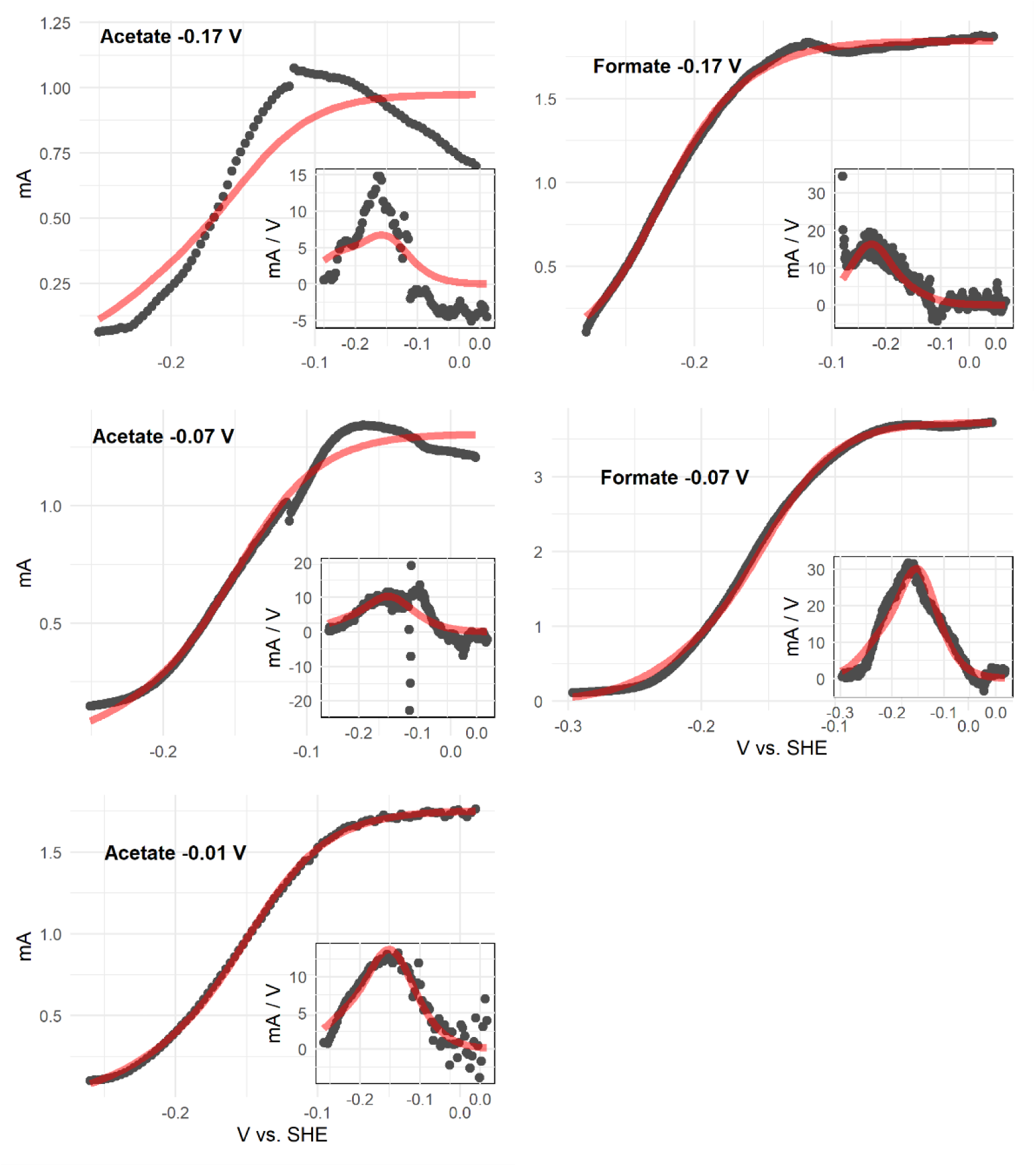
One representative cyclic voltammogram from each condition with its respective Nernst-Monod model fitting. The experimental data is plotted as gray points, and the model fitting is the red line. Labels on each plot describe the electron donor used (acetate or formate) and the fixed anode potential grown at (−0.17 V, −0.07 V, or −0.01 V vs SHE). Inset in each plot is the differentiation of the CV data (slope) plotted in gray and the derivative of the model fit function as a red line. Cyclic voltammograms were collected at a scan rate of 5 mV/s at a biofilm current density between 1-2 A/m^2^ during exponential current increase in order to capture a thin biofilm at a fast enough scan rate to prevent transcriptional adaptation to the shifting potential.

**Table 1:**
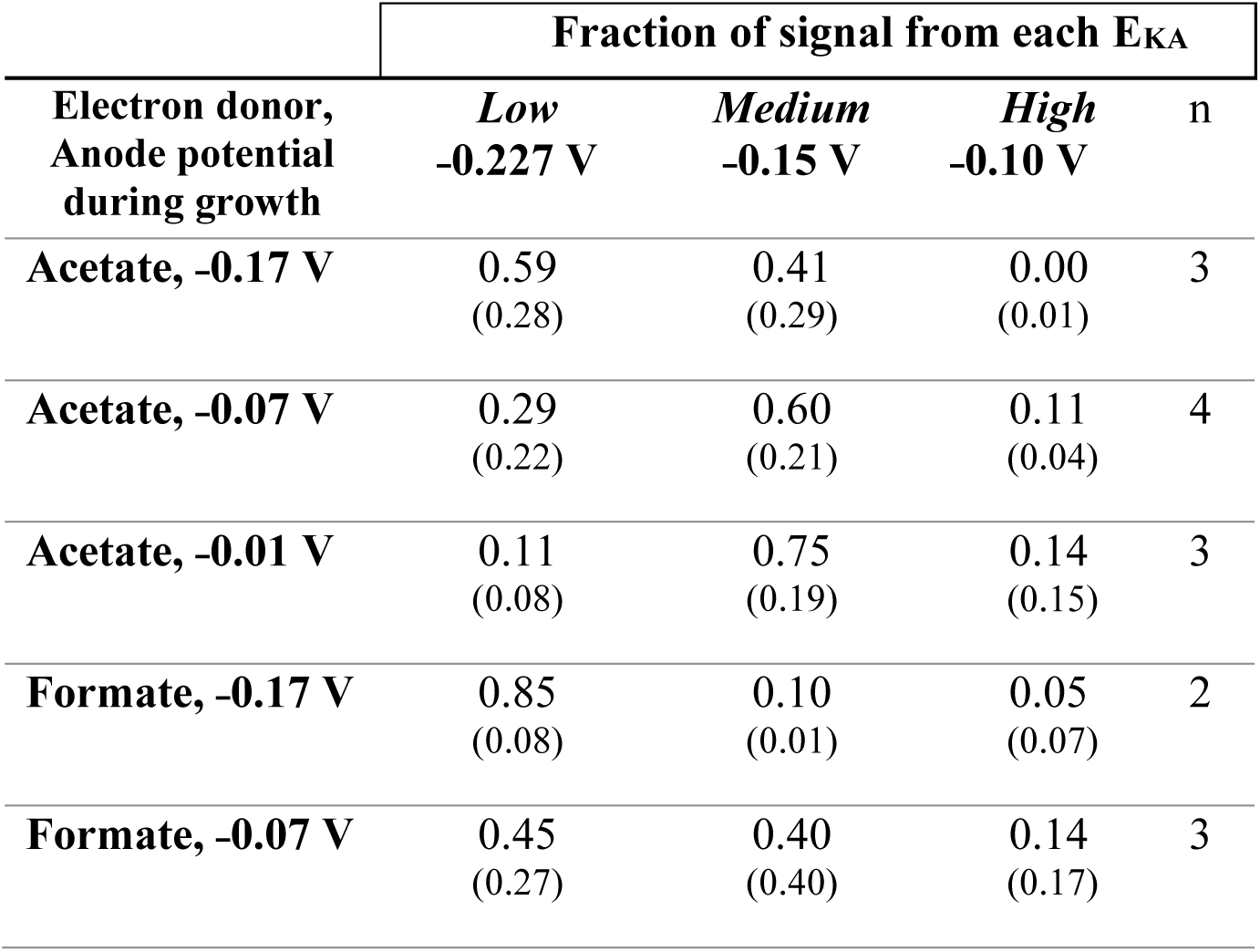
Average fractional contribution of each Nernst-Monod function to the overall CV fit in each condition. A larger fraction represents a stronger contribution of a certain pathway to the overall signal, while a fraction close to zero suggests that a certain pathway does not contribute to the overall signal in its respective growth condition. Calculated as j_max,i_/Σj_max,n_ for each cyclic voltammogram. Numbers in parentheses under each value are the sample standard deviations of each parameter from the number of replicates listed in the ‘n’ column. The “High”, “Medium”, and “Low” E_KA_ values represent the midpoint potentials (vs. SHE) of each Nernst-Monod expression that add up to best fit the experimental data.

| Electron donor,<br>Anode potential<br>during growth | Fraction of signal from each $E_{KA}$ | | | n |
| --- | --- | --- | --- | --- |
|  | *Low*<br>-0.227 V | *Medium*<br>-0.15 V | *High*<br>-0.10 V |  |
| Acetate, -0.17 V | 0.59<br>(0.28) | 0.41<br>(0.29) | 0.00<br>(0.01) | 3 |
| Acetate, -0.07 V | 0.29<br>(0.22) | 0.60<br>(0.21) | 0.11<br>(0.04) | 4 |
| Acetate, -0.01 V | 0.11<br>(0.08) | 0.75<br>(0.19) | 0.14<br>(0.15) | 3 |
| Formate, -0.17 V | 0.85<br>(0.08) | 0.10<br>(0.01) | 0.05<br>(0.07) | 2 |
| Formate, -0.07 V | 0.45<br>(0.27) | 0.40<br>(0.40) | 0.14<br>(0.17) | 3 |

Our cyclic voltammetry fitting with the Nernst-Monod expression indicates that there is an observable difference in which pathways are contributing most to the overall biofilm CV signal (Table 1). The acetate −0.07 V, acetate −0.17 V, and formate −0.07 V conditions had contributions from the Low and Medium signals with little to no contribution from the High pathway. On the other hand, the CVs from the formate −0.17 V condition were overwhelmingly influenced by the Low signal, and CVs from the acetate −0.01 V condition were mainly fitted with the Medium signal. Even though two of the conditions had an anode potential above −0.1 V, none of the CVs had a major contribution from the High signal (Table 1) although higher growth potentials should have a higher contribution of this pathway. The small contributions of the High signal that we observed may be due to the anode potentials we used being lower than the switch point that would trigger expression of the High pathway. Using the information from this model fitting of our CV data, we can discuss differential expression of electron pathway genes in the context of what pathway they may be more strongly associated with.

### Growth conditions affect the overall transcriptome

The main drivers of differential gene expression were the different electron donors and electron acceptors used. The fumarate biofilm and the planktonic fumarate samples are highly differentiated from the anode samples, and among the anode samples there is a clear differentiation between formate and acetate conditions but for acetate there was no clear differentiation based on different anode potentials. All three of the anode biofilms grown on acetate cluster closely together, although there are still some significantly differentially expressed genes. A multi-tab .xlsx file with significantly different genes for all pairwise comparisons is included in the SI. The different anode potentials we used had a small effect on the overall transcriptome of *G. sulfurreducens* grown using acetate, but there is clear separation between the two conditions using formate as the electron donor.

EET pathways in *G. sulfurreducens* are populated with multiheme cytochromes – some known and some unknown. By synthesizing the modeling outputs from our CV data (Table 1) and the differential gene expression of multiheme cytochromes in important pairwise comparisons (Figure 3), we can create associations between cytochromes and electrochemical pathways (Table 2).

**Figure 2:**
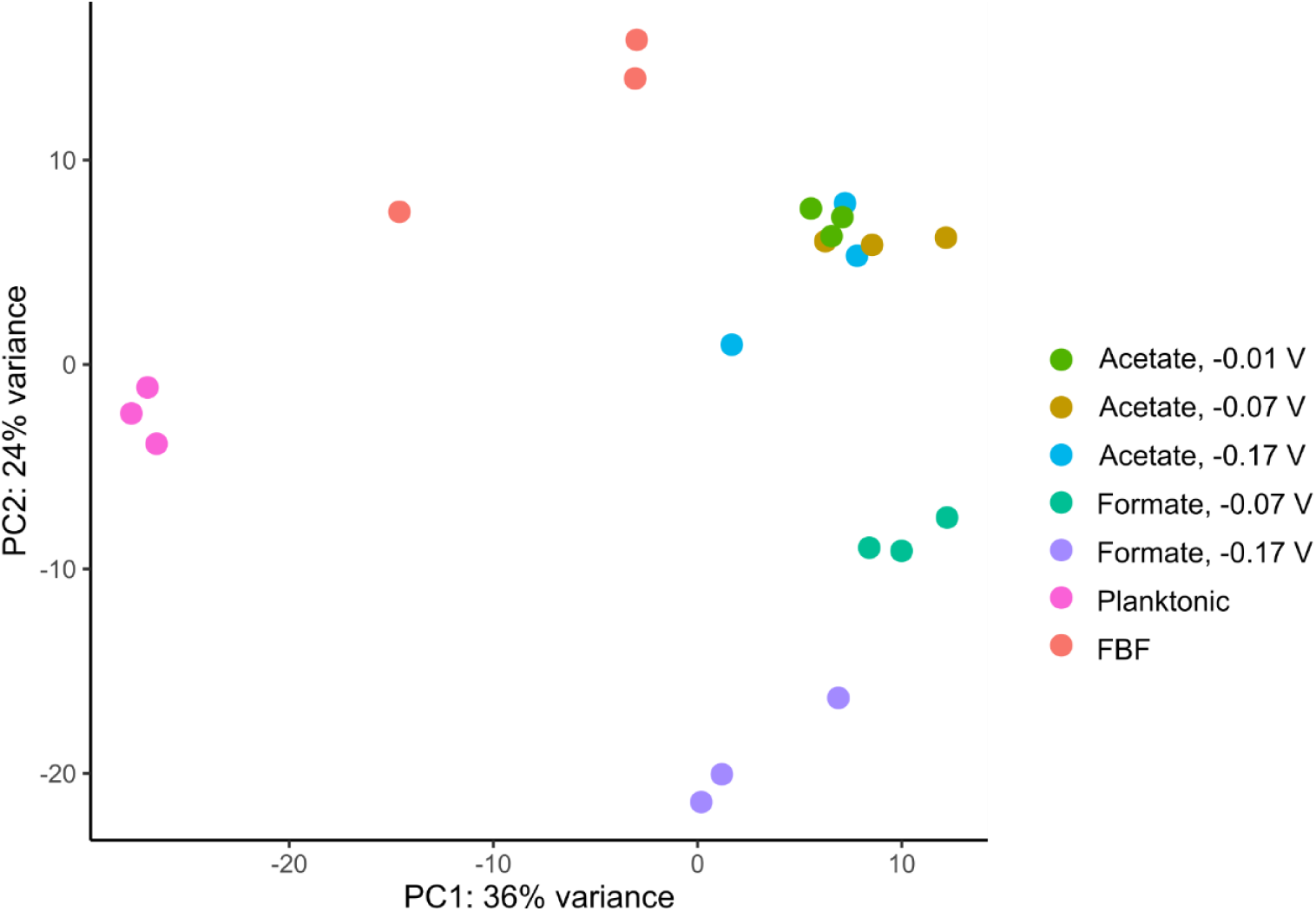
PCA plot of variance in expression between triplicate samples grown under seven different electron donor and acceptor conditions. FBF is a fumarate biofilm condition that was grown on an unconnected graphite rod with fumarate as the electron acceptor.

**Figure 3:**
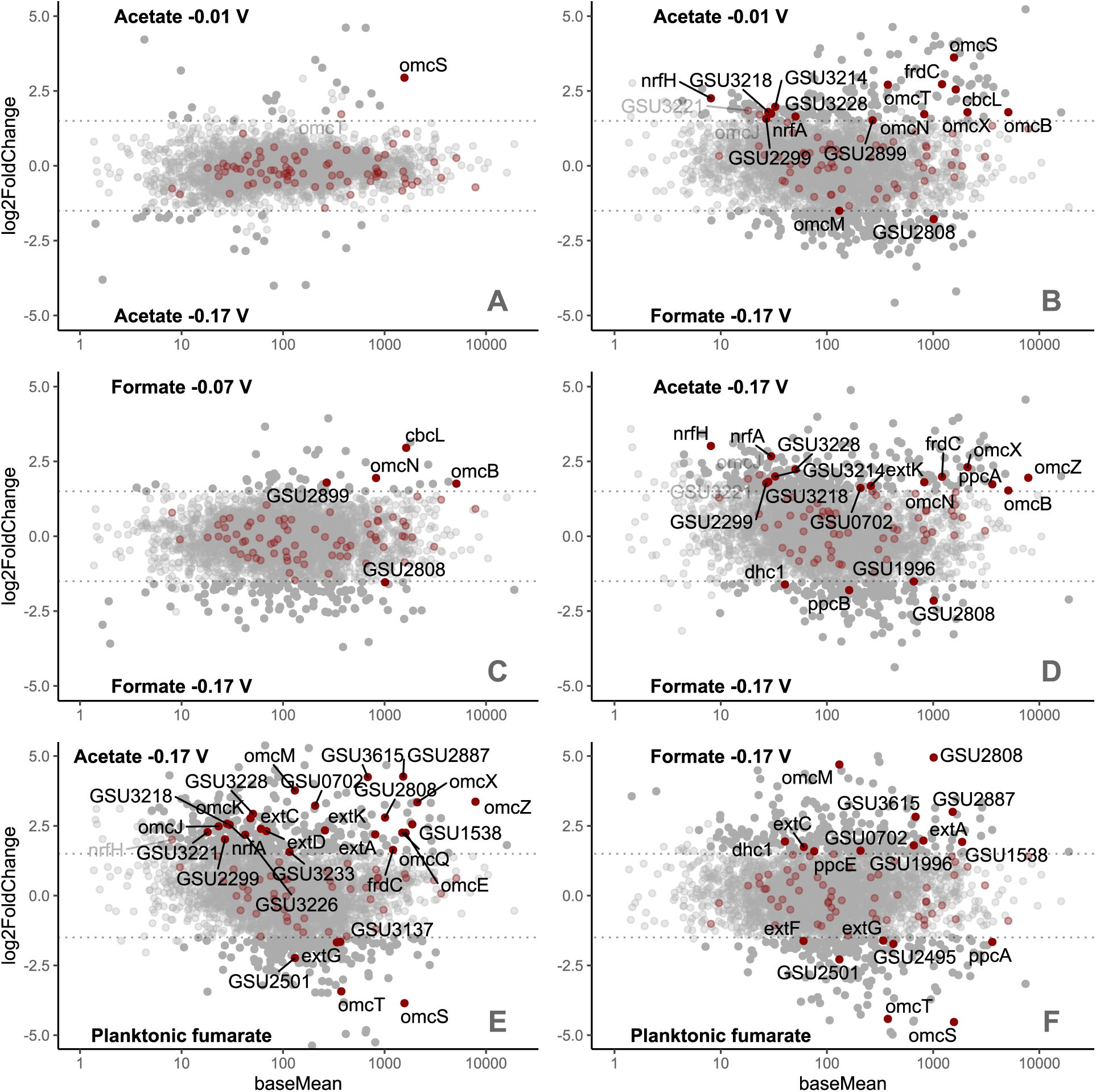
MA plots highlighting interesting pairwise comparisons of differential gene expression with multiheme cytochromes highlighted in red. Labels at top and bottom indicate conditions plotted for comparison. Dotted lines are at ±1.5 log_2_ fold change, and all clearly labeled genes meet an adjusted p-value <0.05. Base mean is calculated using Deseq2’s method for normalizing counts across samples and p values were corrected by the Benjamini Hochberg method. Table 2 lists all the highlighted genes and the associated pathways and cellular locations.

**Table 2:**
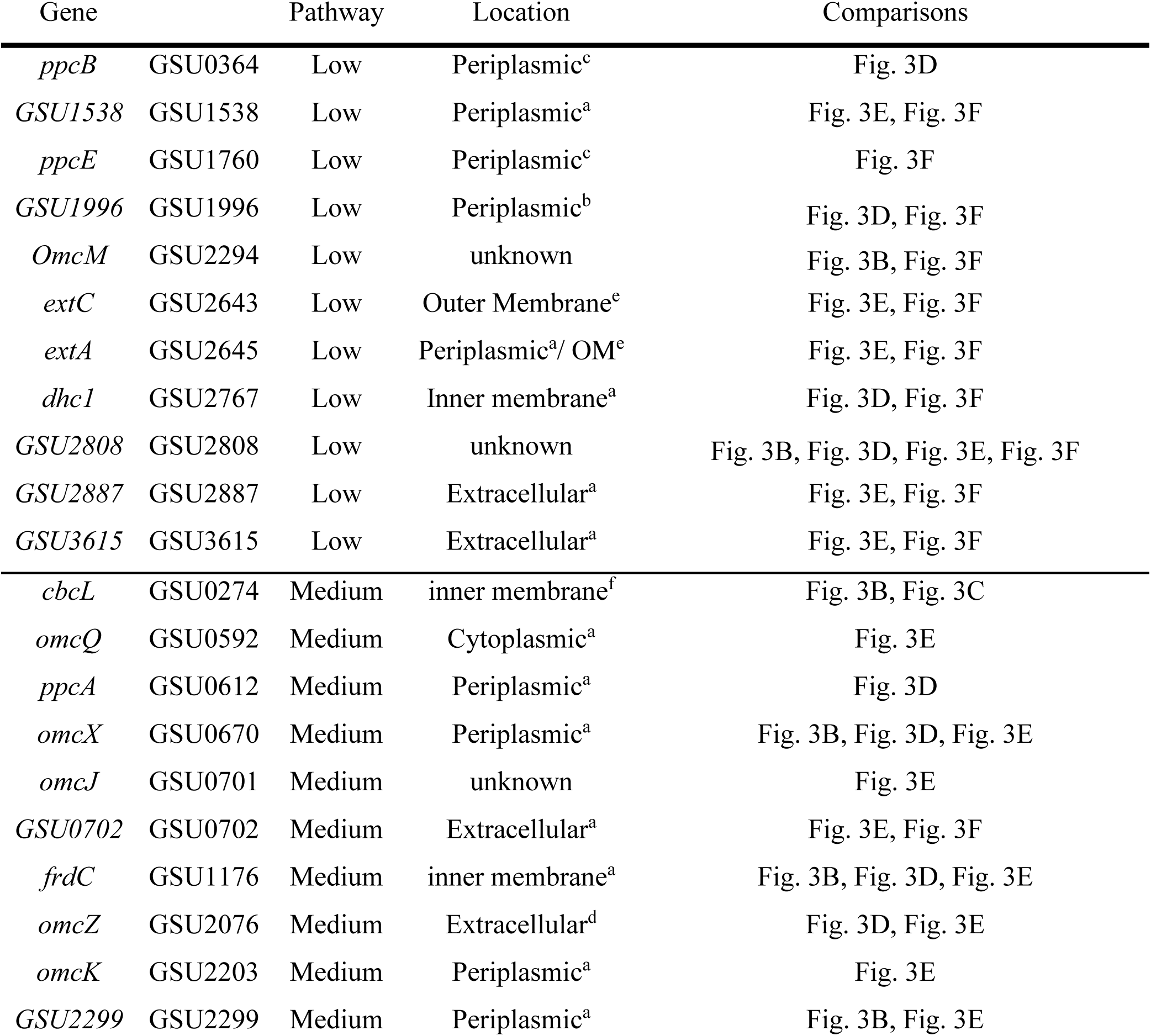
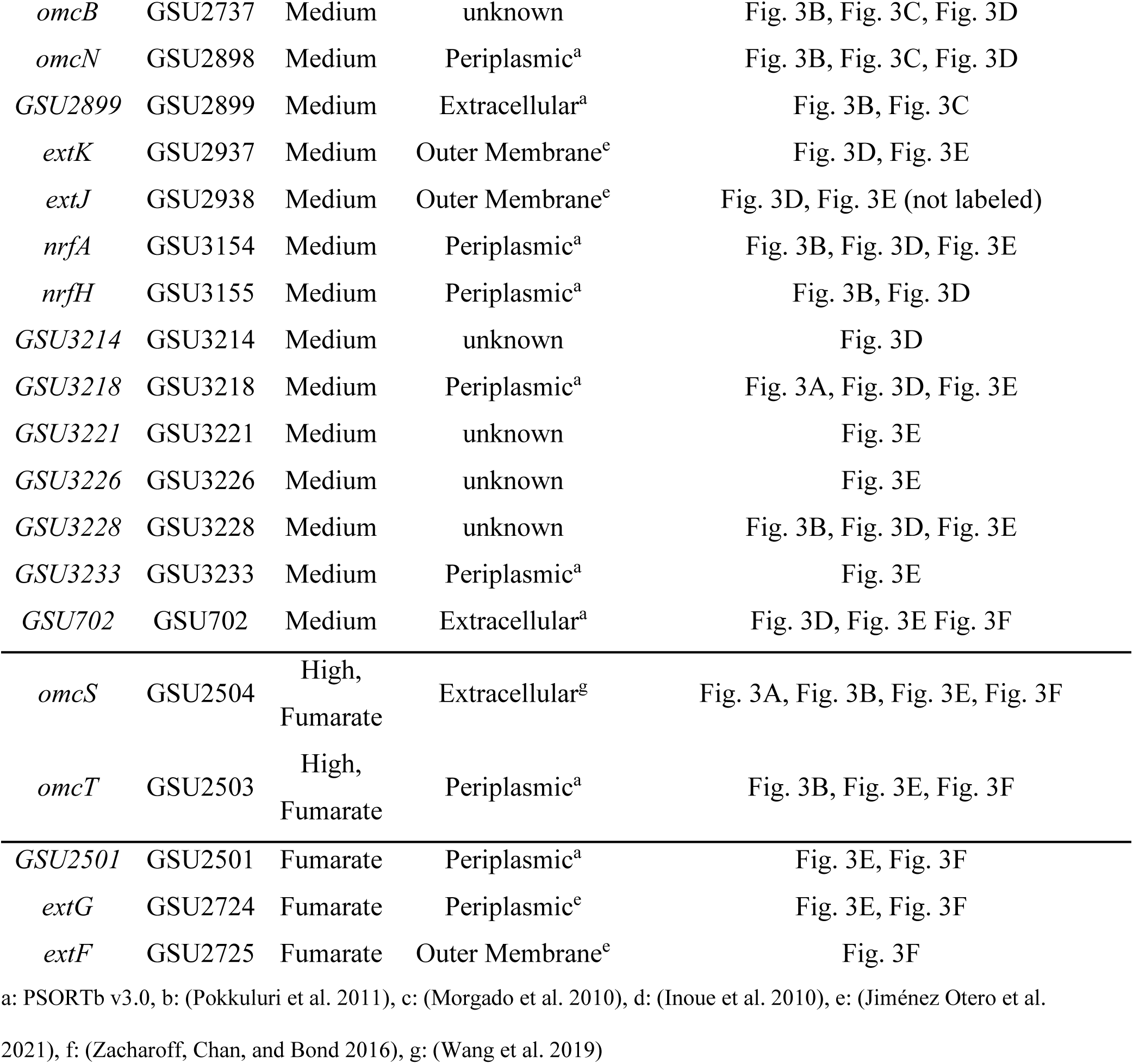
Multiheme cytochromes with significant differential expression and electrochemical data suggesting an association to one of the three EET pathways. These associations were inferred from the differential expression comparisons in Figure 3, and the electrochemical modeling in Table 1. We assume the fumarate samples exhibit primarily the High pathway due to the relatively high redox potential of the fumarate/succinate couple. *extJ* is not a multiheme c-type cytochrome, but we include it here due to its presence in the ExtHIJK cluster.

### Membrane cytochromes can be associated to a specific pathway

Four clusters of outer membrane genes have been shown to be required for EET in *G. sulfurreducens* under certain conditions: ExtABCD, ExtEFG, ExtHIJK, and OmcBC; deleting all four clusters prevents EET respiration (Otero and Chi Ho Chan 2018). Our differential expression data suggests that these clusters may be associated with different electron acceptor potentials. In our data, *extJ* and *extK* were downregulated in the formate −0.17 V biofilm versus the acetate −0.17 V biofilm (log_2_FC=2.6, p_adj_=0.016, log_2_FC=1.7, p_adj_=0.016). The formate −0.17 V biofilm was predominantly represented by the Low pathway in our electrochemical data whereas the acetate −0.17 V biofilm had a strong signal from the Medium pathway. From this, we can infer that the ExtHIJK proteins may contribute to the Medium pathway signal. SHE. Likewise, we infer that ExtEFG may be associated with respiration at the highest potentials due to the downregulation of *extF* and *ExtG* in the formate condition grown at an electrode potential of −0.17 V when compared to the expression in planktonic fumarate sample which we assume represents expression of the pathway associated with the highest potential. The fumarate/succinate redox couple has a standard potential of 0.03 V at pH 7, making it more positive than any other acceptor used in this study.

The expression of the gene *omcB* is downregulated in the formate −0.17 V biofilm relative to the acetate anode biofilms at all three potentials and the formate −0.07 V biofilm, indicating that it may be associated with the Medium pathway. OmcB forms part of a conduit that transfers electrons through the outer membrane and is involved in reduction of Fe(III) citrate and ferrihydrite (Liu et al. 2015). We observed differential expression of OmcB but not OmcC – two paralogs with 65% amino acid similarity. *GSU2808* is another outer membrane cytochrome with the opposite expression pattern to *omcB. GSU2808* was significantly upregulated in the formate −0.17 V biofilm relative to all other anode biofilm conditions indicating an association with the Low pathway. *GSU2808* is a 5 heme cytochrome that has been linked to palladium reduction with acetate as the electron donor (Hernández-Eligio et al. 2020).

We expected to see upregulation of *imcH*. ImcH is the inner membrane cytochrome required for EET at anode potentials above −0.1 V vs. SHE, but there was no significant change among any anode potentials. Yet, *imcH* was a highly expressed cytochrome gene in all conditions, with a base mean of 869 which is in the 94^th^ percentile among all genes. Given the anode potentials we chose, it is possible that our experimental design did not capture the differential expression of *ImcH*. Based on our electrochemical modeling and the differential expression data, none of the conditions we studied were primarily using the High pathway associated to *ImcH* (Table 1,).

CbcL, the inner membrane cytochrome required for potentials below −0.1 V vs. SHE, was significantly downregulated in the formate −0.17 vs. SHE samples compared to the formate −0.07 V samples (log2FC = 3.0, adjusted p = 1.8E−5) but not significantly different between any of the acetate anode biofilm potentials. This observation fits well with the CV data where all conditions except for the −0.17 V formate biofilm had a signal from the Medium pathway. Recent work has identified CbcAB as a crucial protein complex for *G. sulfurreducens* to respire at potentials below −0.2 V vs. SHE (Joshi et al preprint 2021), but we did not detect any differential expression in *cbcA* or *cbcB*, although each gene had a base mean of counts in the 97^th^ and 95^th^ percentile, respectively. Apart from these known inter membrane proteins, *dhc1* was differentially expressed with an association to the Low potential condition (Table 2). This gene encodes for a unique diheme cytochrome of unknown function with a predicted inner membrane localization.

### Extracellular cytochromes network adapts to redox conditions

Filaments made of OmcS or OmcZ units have been implicated as a critical component of long distance EET (Wang et al. 2019; Yalcin et al. 2020). We observed that *omcS* was significantly upregulated in the fumarate conditions relative to all anode biofilms except for the acetate −0.07 V vs. SHE condition. *G. sulfurreducens* may use OmcS for EET only at higher anode potentials, above ∼ (−0.1) V vs. SHE. Fumarate, with a standard reduction potential of 0.03 V vs. SHE, appears to be stimulating the expression of the High pathway. To our knowledge, previous studies on OmcS filaments have all used either fumarate or an electrode poised above 0 V vs. SHE as the electron acceptor. Of the “*omc*” genes, the most likely candidate for replacing the function of OmcS at lower potentials based on our data is *omcZ*. The OmcZ protein was recently shown to form nanowires with a high conductivity in *G. sulfurreducens* (Yalcin et al. 2020) and has been shown to be preferentially expressed at low potentials (Peng and Zhang 2017). Our data supports that the differential expression of *omcS* vs. *omcZ* appears to be at least partially controlled by electron acceptor potential with *omcZ* being important when the Medium pathway is detected. Interestingly, we did not observe an upregulation of *omcZ* in the Formate −0.17 V condition relative to planktonic fumarate despite the down regulation of *omcS* (Figure 3B and 3F), suggesting that there may be more than just *omcS* and *omcZ* responsible for the pathway-dependent switch in the extracellular part of the pathway. Other cytochromes with predicted extracellular localization (GSU2887, GSU3615) were more abundantly expressed under the Low Pathway conditions, but the function of these cytochromes is yet to be elucidated (Kim et al. 2005; Leang et al. 2005; Peng et al. 2016)

### Differential expression of Ppc proteins may maximize respiratory efficiency

Five periplasmic triheme cytochromes (PpcA-E) have been identified and characterized in *G. sulfurreducens* (Lloyd et al. 2003). PpcA was first characterized and shown that its deletion affects Fe(III) and U(VI) reduction, but not fumarate reduction. PpcB-D were initially identified due to their high similarity to PpcA (75-100%), while predictions confirmed their periplasmic location (Shelobolina et al. 2007). Most studies have suggested the role of these cytochromes is to transfer electrons collected at the inner membrane to outer-membrane cytochromes (Lloyd et al. 2003; Morgado et al. 2010; Shelobolina et al. 2007). Interestingly, further studies have confirmed that PpcA and PpcD perform a proton-coupled electron transfer; which leads to the possibility that these periplasmic cytochromes are associated to ATP production (Morgado et al. 2010). It has also been confirmed that electron transfer by PpcB and PpcE are not proton coupled, suggesting a different cellular function compared to PpcA/PpcD (Silva, Portela, and Salgueiro 2021).

Our observed gene expression for Formate at −0.17 V vs SHE, which has primarily the Low pathway evident, shows an increased expression of PpcB (Fig. 3D) and PpcE (Fig. 3F) over PpcA compared to conditions where the Medium pathway is active. Our study associates the use of PpcB and PpcE to the Low pathway. On the other hand, *ppcA* seems to be highly expressed in all other samples, this suggests its utilization for the Medium and/or High pathways. Given that the lower potential pathway is associated with lower electron-transfer energy available in *G. sulfurreducens* metabolism, there is limited energy for proton pumping and ATP production.

The use of PpcB/PpcE could be an approach to decrease ATP production when the energy gradient is limiting, while PpcA is used when more energy is available. Thus, our results are consistent with the hypotheses that PpcA/PpcD are associated with ATP production (Morgado et al. 2010; Silva, Portela, and Salgueiro 2021). Analyses of these periplasmic cytochromes have also discussed differences in working potentials, which could indicate different electron acceptors (Morgado et al. 2010). The PpcA-E proteins function as periplasmic electron carriers in *G. sulfurreducens’* EET, so a change in the distribution of the different Ppc proteins could allow *G. sulfurreducens* to maintain efficiency as external redox conditions change.

We also observed differential expression of several large cytochromes, each with at least a dozen hemes. *GSU3218* (15 hemes), *GSU0702* (35 hemes), and *omcN* (34 hemes) are upregulated in the acetate −0.17 V condition relative to the formate −0.17 V condition, and therefore we associate them with the Medium pathway (Table 2). *GSU0702* has been previously associated with respiration of electrodes at lower potentials as it was upregulated in *G. sulfurreducens* grown at −0.25 V vs. SHE compared to +0.2 V (Peng et al. 2016). Deleting *omcN* does not impact the reduction of soluble or insoluble electron acceptors in one study (Aklujkar et al. 2013). The 27-heme cytochrome gene *GSU2887* is associated with the Low pathway. The proteins encoded by *GSU2887* and *GSU0702* are both predicted to be extracellular by PSORTb 3.0, but it is not clear what the function of such large cytochromes would be outside the cell.

### Formate metabolism suppresses the TCA cycle

When acetate is available, *Geobacter* uses NADH generated from the TCA cycle to pump protons across the membrane using NADH dehydrogenase for energy conservation. When formate is the only electron donor available, *Geobacter* suppresses the TCA cycle and shifts to an alternate metabolism. There were 80 genes differentially expressed between *G. sulfurreducens* anode biofilms grown with the same −0.07 V vs. SHE potential with either acetate or formate as the electron acceptor. Ten genes encoding for subunits of the NADH dehydrogenase complex were significantly downregulated in the formate condition. Every gene encoding for an enzyme in the TCA cycle except for *sucA*, 2-oxoglutarate dehydrogenase, was downregulated in the formate samples. In the formate condition, we observed the upregulation of 15 genes including *fdhD* (formate dehydrogenase) (Supplemental spreadsheet). The downregulation of NADH dehydrogenase (*nuo* genes) and the upregulation of formate dehydrogenase that we observe in formate biofilms indicate that formate dehydrogenase is replacing NADH dehydrogenase as the primary electron carrier in the membrane electron transfer chain. There is no evidence that *Geobacter*’s formate dehydrogenase can pump protons in the same way as NADH dehydrogenase.

Formate may stimulate a similar response as hydrogen in *Geobacter. hgtR*, the hydrogen-dependent growth transcriptional regulator, is a critical protein for shifting metabolism to use hydrogen as the electron donor (Ueki and Lovley 2009). Joshi, Chan, and Bond 2021 also found that *hgtR* is upregulated in a mutant *G. sulfurreducens* where an upstream regulator of CbcAB is deleted (Δ*bccR*), potentially because of thermodynamic limitations on respiration. We observed upregulation of *hgtR* in both formate biofilms relative to all acetate conditions except for the acetate −0.01 V biofilm. Although we did observe downregulation of the TCA cycle in formate conditions, the fact that *hgtR* was also relatively highly expressed in the −0.01 V acetate biofilm suggests that it is not solely related to hydrogen/formate metabolism, but instead a thermodynamic response in *G. sulfurreducens*. It is also possible that small amounts of hydrogen were produced in our single-chamber reactors, and this activated *hgtR*, but in that case we would not expect to see differential expression as a function of potential or electron donor.

We initially chose to study conditions where fumarate was the electron acceptor to capture the transcriptome of cells that did not need to use EET. Instead, we observed the expression of many of the EET-associated proteins in the fumarate conditions. Our data support other work that has shown that *G. sulfurreducens* is not optimized to grow with fumarate. OmcS nanowires have been isolated from fumarate cultures of *G. sulfurreducens* (Wang et al. 2019), indicating that fumarate cultures produce the machinery of the High potential EET pathway. *G. sulfurreducens* also prefers Fe(III) as an electron acceptor when both Fe(III) and fumarate are available despite fumarate providing more energy (Esteve-Núñez, Núñez, and Lovley 2004). *G. sulfurreducens* uses a reversible enzyme complex, FrdCAB for both fumarate reduction during fumarate respiration and succinate dehydrogenation as part of the TCA cycle (Butler et al. 2006). In our data *frdC* was differentially expressed, although it was not correlated with fumarate utilization. Rather, fumarate reductase was regulated along with the other elements of the TCA cycle. *frdC* was downregulated in the formate conditions relative to the acetate biofilms which is consistent with the downregulation of the other TCA cycle enzymes. We did, however, observe an increased expression of *dcuB* in fumarate conditions. DcuB is the fumarate transporter in *G. sulfurreducens* (Butler et al. 2006).

## Conclusion

Due to its complicated electron pathways, fully understanding EET in *G. sulfurreducens* will require synthesizing data from many techniques, and a large body of literature. In this study, we combined electrochemical CV fittings using Nernst-Monod modeling and RNASeq to identify electrochemical conditions that change the expression of genes associated with different pathways. We have identified a number of genes that show an association to different respiratory pathways in *G. sulfurreducens*, and this technique could be expanded in further studies to answer more unresolved questions about the metabolism of *G. sulfurreducens* and other electroactive organisms.

Over decades of research in *G. sulfurreducens*, expression studies have often assumed an anode, often poised at high potentials, as a fixed condition for growth; only recently have studies started to recognize the importance of shifting anode potentials in its expression and electrochemical behavior (Levar et al. 2017; Peng et al. 2016). Our results suggest that a small shift in the anode potential (100-160 mV in our study) cause significant changes in the respiratory pathway of *G. sulfurreducens*. While this is a small potential change, it constitutes a significant increase (87-139%) in available energy for growth using acetate as electron donor (E^0’^= − 285 mV vs SHE). It seems that changes along the full electron path, from the inner membrane to the extracellular matrix, occur because of a change in potential. In Figure 4, we hypothesize an updated model of EET in *G. sulfurreducens*. Trifurcation of electrons occurs at the inner membrane as previously reported (Joshi, Chan, and Bond 2021; Levar et al. 2017). In order to maintain efficient energy conservation, many redox proteins along the EET pathway should change to create a cascading potential gradient from the inner membrane to the anode. Based on our data, we propose in Figure 4 some of the electron carriers in each of the three pathways, with changes at the periplasmic, outer membrane, and extracellular portions of the electron transfer chain. We anticipate that future studies will associate more electron carriers with each pathway and complete the model.

**Figure 4:**
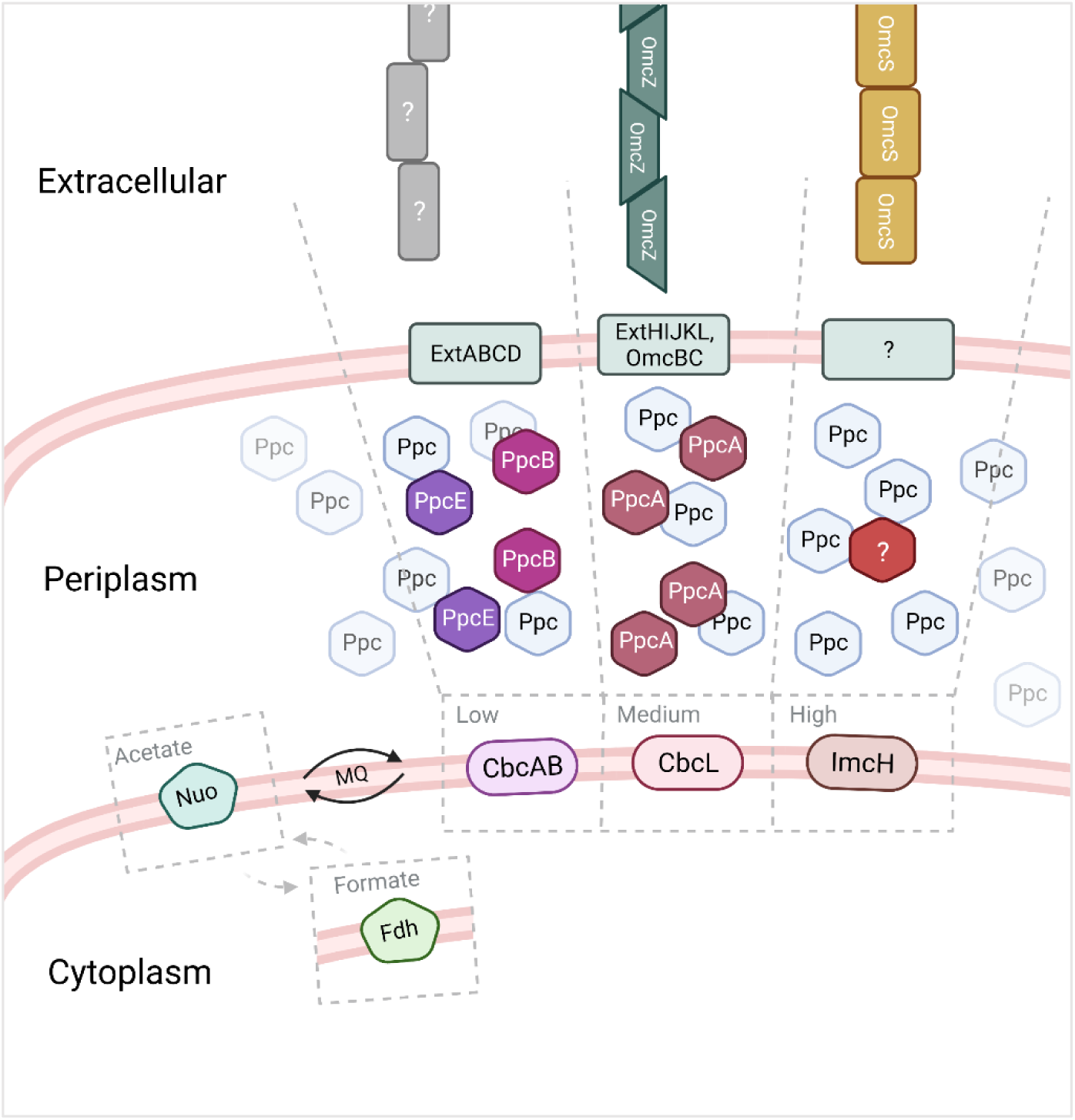
Our hypothesized schematic of the EET process in *G. sulfurreducens* and how it is differentiated based on electron acceptor potential. At the inner membrane NADH dehydrogenase (Nuo) reduces menaquinone (MQ), or in the case of formate metabolism Nuo is downregulated and formate dehydrogenase (Fdh) initializes the inner membrane electron transfer chain. At the inner membrane, electrons enter one of three EET pathways. In the Low pathway, CbcAB (Joshi, Chan, and Bond 2021) oxidizes menaquinol, the periplasmic electron transfer favors PpcB and PpcE, and the ExtABCD complex is involved at the outer membrane. In the Medium pathway, CbcL transfers electrons to PpcA in the periplasm and then to ExtHIJKL and OmcBC and OmcZ at the outer membrane. Our study did not capture the High pathway well, but we do know it uses ImcH (Levar et al. 2017) at the inner membrane and OmcS as an extracellular electron carrier. This diagram is not a complete explanation of all the EET pathways in *G. sulfurreducens* but rather a way to highlight some of the associations we have discovered. Created with BioRender.com.

**Figure 5:**
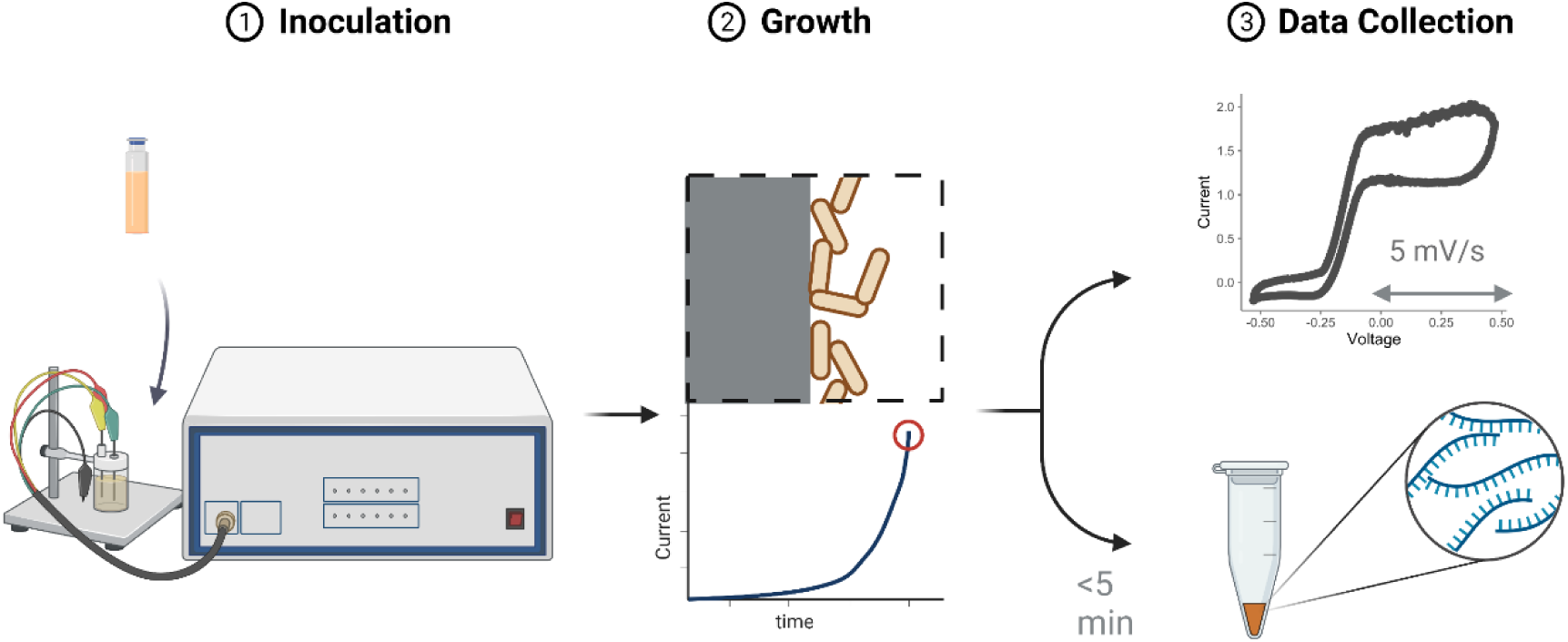
1) *G. sulfurreducens* is inoculated into a microbial electrochemical cell with a graphite electrode acting as the electron acceptor for growth using a potentiostat to apply a potential. 2) Growth is monitored via chronoamperrometry until the bacteria produce 1-2 A/m^2^ during exponential growth, indicating a thin and active biofilm. 3) At this stage, we perform cyclic voltammetry on the biofilm, or the biofilm is collected, and RNA is extracted for transcriptomics. Created with BioRender.com.

Since the respiratory pathways in *G. sulfurreducens* involve a complex network of cytochromes extending to the extracellular space, changing this pathway due to a shift in electron acceptor conditions (e.g., change in anode or metal oxide potential) would require a significant energy investment. Thus, our results imply some limitations in respiratory efficiencies as redox conditions change and add more context to the complicated story of *G. sulfurreducens*.

## Methods

### Growth

*Geobacter sulfurreducens* PCA was grown under 7 different conditions. “Planktonic” samples used ATCC 1957 medium in sealed anaerobic test tubes and extracted 48 hours after inoculation. Anode biofilm samples used a modified ATCC 1957 medium without sodium fumarate in a single chamber microbial electrochemical cell with a graphite rod electrode (600-800mm^2^) serving as an anode. The reactor volume was 100mL with an Ag/AgCl reference electrode (BASi, Indiana USA) and a stainless-steel wire as cathode. Acetate anode biofilms were poised at three different potentials, −0.07 V vs. SHE, −0.17 V vs. SHE, and −0.01 V vs. SHE. Formate anode biofilms were poised at two potentials – −0.07 V vs. SHE, and −0.17 V vs. SHE. We used a conversion of −0.27 V vs. Ag/AgCl is 0 V vs. SHE. Anode biofilm growth was monitored with a BioLogic VMP3 potentiostat (Tennessee, USA) and samples were collected during exponential current growth phase in the range of 1-3 A/m^2^ to capture a thin biofilm. Fumarate biofilm samples used ATCC 1957 medium in a continuously mixed flow-through reactor identical to the anode biofilm reactor with a hydraulic retention time less than the reported doubling time for *G. sulfurreducens* and with the cell in open circuit. Planktonic samples were collected by centrifugation, and biofilms were collected by scraping. Seven conditions were sampled in biological triplicate for a total of twenty-one samples.

### RNA processing

RNA was extracted from all samples with a QIAGEN PowerMicrobiome RNA extraction kit (Germany). The extraction process began within 5 minutes of disturbing each culture. RNA quality and quantity were assessed with a Nanodrop spectrophotometer and an Agilent Bioanalyzer 2100. Ribosomal RNA contamination was reduced with the MICROBExpress Bacterial mRNA enrichment kit (ThermoFisher, Massachusetts USA). Illumina (California USA) sequencing libraries were prepared with KAPA Biosystems Hyperprep RNA (Massachusetts USA) before pooling and sequencing on an Illumina Nextseq 500 2×150 module. Sequencing was performed at the OKED Genomics Core at Arizona State University.

### Differential Expression

Sequence quality was assessed with FastQC (Andrews 2010). After demultiplexing, reads were trimmed with trimmomatic (Bolger, Lohse, and Usadel 2014) to remove low quality read segments. The majority of ribosomal reads were removed by alignment to reference sequences in Bowtie2 Filtered reads were aligned to the *G. sulfurreducens* PCA RefSeq genome using Bowtie2 (Langmead and Salzberg 2012), and only paired alignments were kept. Alignment files were transformed for analysis in R with samtools (Li et al. 2009). In R, aligned reads were mapped to the NCBI RefSeq assembly for *G. sulfurreducens*. Ribosomal RNA-mapped reads were removed before differential expression analysis. Differential gene expression was analyzed with the R package DESeq2 (Love, Huber, and Anders 2014). Ribosomal protein genes are removed from data presented here. Transcriptomic data and metadata are hosted in the NCBI Gene Expression Omnibus database under accession number GSE200066.

### Electrochemistry

Anode biofilms used for cyclic voltammetry were grown under similar conditions to the samples used for RNA extraction. We used a Bio-Logic VMP3 potentiostat for all chronoamperometry and cyclic voltammetry. Cyclic voltammograms were recorded with a scan rate of 5 mV/s from −0.5 V vs. SHE to +0.5 V vs SHE. A total of 3 consecutive scans were performed for each biofilm and the second scan was used for fittings. Fitting to Nernst-Monod curves to the data by optimizing each j_max_ was performed in using the nl2sol non-linear least squares regression from the R package stats (R 3.6.3, 2021).

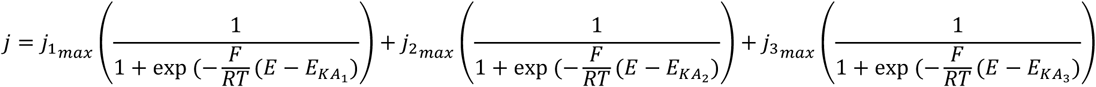

Equation 1: Three-part Nernst-Monod expression to model the *G. sulfurreducens* biofilm cyclic voltammograms. F is Faraday’s constant (96,485 C/mol), R is ideal gas law constant (−8.314 J/mol K, T is temperature (K), and E is the anode potential (V) as measured by the potentiostat. j is current density (A/m^2^). In the model, each j_i,max_ was a fitting parameter.

## Acknowledgements

The funding for this work was provided by the Office of Naval Research (ONR awards N0014-15-1-2702 and N0014-20-1-2269).

## Notes

https://www.ncbi.nlm.nih.gov/geo/query/acc.cgi?acc=GSE200066

### Competing Interest Statement

The authors have declared no competing interest.

## References

Aklujkar, M. et al. 2013. “Proteins Involved in Electron Transfer to Fe(III) and Mn(IV) Oxides by Geobacter Sulfurreducens and Geobacter Uraniireducens.” Microbiology (United Kingdom) 159(PART3): 515–35.

Andrews, S. 2010. “FastQC: A Quality Control Tool for High Throughput Sequence Data [Online].” http://www.bioinformatics.babraham.ac.uk/projects/fastqc/.

Bolger, Anthony M., Marc Lohse, and Bjoern Usadel. 2014. “Trimmomatic: A Flexible Trimmer for Illumina Sequence Data.” Bioinformatics 30(15): 2114–20.

Bonanni, Pablo Sebastián, Diego Massazza, and Juan Pablo Busalmen. 2013. “Stepping Stones in the Electron Transport from Cells to Electrodes in Geobacter Sulfurreducens Biofilms.” Physical Chemistry Chemical Physics 15(25): 10300–306.

Bond, Daniel R., and Derek R. Lovley. 2003. “Electricity Production by Geobacter Sulfurreducens Attached to Electrodes.” AEM 69(3): 1548–55.

Butler, Jessica E et al. 2006. “Genetic Characterization of a Single Bifunctional Enzyme for Fumarate Reduction and Succinate Oxidation in Geobacter Sulfurreducens and Engineering of Fumarate Reduction in Geobacter Metallireducens.” 188(2): 450–55.

Caccavo, Frank et al. 1994. “Geobacter Sulfurreducens Sp. Nov., a Hydrogen-and Acetate-Oxidizing Dissimilatory Metal-Reducing Microorganism.” Applied and environmental microbiology 60(10): 3752–59.

Clarke, T. A. et al. 2011. “Structure of a Bacterial Cell Surface Decaheme Electron Conduit.” Proceedings of the National Academy of Sciences 108(23): 9384–89. http://www.pnas.org/cgi/doi/10.1073/pnas.1017200108.

Ding, Yan Huai R et al. 2008. “Proteome of Geobacter Sulfurreducens Grown with Fe(III) Oxide or Fe(III) Citrate as the Electron Acceptor.” Biochimica et Biophysica Acta - Proteins and Proteomics 1784(12): 1935–41. http://dx.doi.org/10.1016/j.bbapap.2008.06.011.

Esteve-Núñez, Abraham, Cinthia Núñez, and Derek R. Lovley. 2004. “Preferential Reduction of Fe(III) over Fumarate by Geobacter Sulfurreducens.” Journal of Bacteriology 186(9): 2897–99.

Gao, Yifan et al. 2018. “Urine-Powered Synergy of Nutrient Recovery and Urine Purification in a Microbial Electrochemical System.” Environmental Science: Water Research and Technology 4(10): 1427–38.

Hernández-Eligio, Alberto et al. 2020. “Global Transcriptional Analysis of Geobacter Sulfurreducens under Palladium Reducing Conditions Reveals New Key Cytochromes Involved.” Applied Microbiology and Biotechnology 104(9): 4059–69.

Holmes, Dawn E. et al. 2006. “Microarray and Genetic Analysis of Electron Transfer to Electrodes in Geobacter Sulfurreducens.” Environmental Microbiology 8(10): 1805–15.

Inoue, Kengo et al. 2010. “Purification and Characterization of OmcZ, an Outer-Surface, Octaheme c -Type Cytochrome Essential for Optimal Current Production by Geobacter Sulfurreducens ? †.” 76(12): 3999–4007.

Jiménez Otero, Fernanda et al. 2021. “Evidence of a Streamlined Extracellular Electron Transfer Pathway from Biofilm Structure, Metabolic Stratification, and Long-Range Electron Transfer Parameters.” Applied and Environmental Microbiology 87(17).

Joshi, Komal, Chi Ho Chan, and Daniel R Bond. 2021. “Geobacter Sulfurreducens Inner Membrane Cytochrome CbcBA Controls Electron Transfer and Growth Yield near the Energetic Limit of Respiration.” Molecular Microbiology 116(4): 1124–39. https://doi.org/10.1111/mmi.14801.

Kim, Byoung-chan Chan et al. 2005. “OmcF, a Putative c -Type Monoheme Outer Membrane Cytochrome Required for the Expression of Other Outer Membrane Cytochromes in Geobacter Sulfurreducens.” Journal of bacteriology 187(13): 4505–13.

Langmead, Ben, and Steven L. Salzberg. 2012. “Fast Gapped-Read Alignment with Bowtie 2.” Nature Methods 9(4): 357–59.

Leang, Ching et al. 2005. “Adaptation to Disruption of the Electron Transfer Pathway for Fe(III) Reduction in Geobacter Sulfurreducens.” Journal of Bacteriology 187(17): 5918–26.

Levar, Caleb E et al. 2017. “Redox Potential as a Master Variable Controlling Pathways of Metal Reduction by Geobacter Sulfurreducens.” 11(3): 741–52. http://dx.doi.org/10.1038/ismej.2016.146.

Levar, Caleb E, Chi Ho Chan, Misha G Mehta-kolte, and Daniel R Bond. 2014. “An Inner Membrane Cytochrome Required Only for Reduction Of.” mBio 5(6): 02034–14.

Li, Heng et al. 2009. “The Sequence Alignment/Map Format and SAMtools.” Bioinformatics 25(16): 2078–79.

Liu, Yimo, James K. Fredrickson, John M. Zachara, and Liang Shi. 2015. “Direct Involvement of OmbB, OmaB, and OmcB Genes in Extracellular Reduction of Fe(III) by Geobacter Sulfurreducens PCA.” Frontiers in Microbiology 6(OCT): 1–8.

Lloyd, Jon R. et al. 2003. “Biochemical and Genetic Characterization of PpcA, a Periplasmic c-Type Cytochrome in Geobacter Sulfurreducens.” Biochemical Journal 369(1): 153–61.

Love, Michael I., Wolfgang Huber, and Simon Anders. 2014. “Moderated Estimation of Fold Change and Dispersion for RNA-Seq Data with DESeq2.” Genome Biology 15(12): 1–21.

Malvankar, Nikhil S. et al. 2011. “Tunable Metallic-like Conductivity in Microbial Nanowire Networks.” Nature Nanotechnology 6(9): 573–79.

Morgado, Leonor et al. 2010. “Thermodynamic Characterization of a Triheme Cytochrome Family From Geobacter Sulfurreducens Reveals Mechanistic And Functional Diversity.” Biophysical Journal 99(1): 293–301. http://dx.doi.org/10.1016/j.bpj.2010.04.017.

Nevin, Kelly P. et al. 2009. “Anode Biofilm Transcriptomics Reveals Outer Surface Components Essential for High Density Current Production in Geobacter Sulfurreducens Fuel Cells.” PLoS ONE 4(5).

Otero, Fernanda Jiménez, and Daniel R. Bond Chi Ho Chan. 2018. “Identification of Different Putative Outer Membrane Electron Conduits Necessary for Fe(III) Citrate, Fe(III) Oxide, Mn(IV) Oxide, or Electrode Reduction by Geobacter Sulfurreducens.” Journal of Bacteriology 200(19): 1–20.

Pat-Espadas, Aurora M., Elías Razo-Flores, J. Rene Rangel-Mendez, and Francisco J. Cervantes. 2013. “Reduction of Palladium and Production of Nano-Catalyst by Geobacter Sulfurreducens.” Applied Microbiology and Biotechnology 97(21): 9553–60.

Peng, Luo et al. 2016. “Geobacter Sulfurreducens Adapts to Low Electrode Potential for Extracellular Electron Transfer.” Electrochimica Acta 191: 743–49. http://dx.doi.org/10.1016/j.electacta.2016.01.033.

Peng, Luo, and Yong Zhang. 2017. “Cytochrome OmcZ Is Essential for the Current Generation by Geobacter Sulfurreducens under Low Electrode Potential.” Electrochimica Acta 228: 447–52. http://dx.doi.org/10.1016/j.electacta.2017.01.091.

Pessanha, Miguel et al. 2006. “Thermodynamic Characterization of Triheme Cytochrome PpcA from Geobacter Sulfurreducens : Evidence for a Role Played in e - / H + Energy Transduction.” Biochemistry 45: 13910–17.

Pokkuluri, P. R. et al. 2011. “Structure of a Novel Dodecaheme Cytochrome c from Geobacter Sulfurreducens Reveals an Extended 12nm Protein with Interacting Hemes.” Journal of Structural Biology 174(1): 223–33. http://dx.doi.org/10.1016/j.jsb.2010.11.022.

Richter, Hanno et al. 2009. “Cyclic Voltammetry of Biofilms of Wild Type and Mutant Geobacter Sulfurreducens on Fuel Cell Anodes Indicates Possible Roles of OmcB, OmcZ, Type IV Pili, and Protons in Extracellular Electron Transfer.” Energy and Environmental Science 2(5): 506–16.

Salgueiro, Carlos A. et al. 2022. “From Iron to Bacterial Electroconductive Filaments: Exploring Cytochrome Diversity Using Geobacter Bacteria.” Coordination Chemistry Reviews 452: 214284. https://doi.org/10.1016/j.ccr.2021.214284.

Santos, Telma C. et al. 2015. “Diving into the Redox Properties of Geobacter Sulfurreducens Cytochromes: A Model for Extracellular Electron Transfer.” Dalton Transactions 44(20): 9335–44. http://www.ncbi.nlm.nih.gov/pubmed/25906375.

Shelobolina, Evgenya S. et al. 2007. “Importance of C-Type Cytochromes for U(VI) Reduction by Geobacter Sulfurreducens.” BMC Microbiology 7(Vi): 1–15.

Silva, Marta A., Pilar C. Portela, and Carlos A. Salgueiro. 2021. “Rational Design of Electron/Proton Transfer Mechanisms in the Exoelectrogenic Bacteria Geobacter Sulfurreducens.” Biochemical Journal 478(14): 2871–87.

Torres, César I. et al. 2010. “A Kinetic Perspective on Extracellular Electron Transfer by Anode-Respiring Bacteria.” FEMS Microbiology Reviews 34(1): 3–17.

Torres, César I, Andrew Kato Marcus, Prathap Parameswaram, and Bruce E Rittmann. 2008. “Kinetic Experiments for Evaluating the Nernst-Monod Model for Anode-Respiring Bacteria (ARB) in a Biofilm Anode.” Environmental Science & Technology 42(17): 6593–97. 10.1021/es800970w.

Ueki, Toshiyuki, and Derek R. Lovley. 2009. “Genome-Wide Gene Regulation of Biosynthesis and Energy Generation by a Novel Transcriptional Repressor in Geobacter Species.” Nucleic Acids Research 38(3): 810–21.

Wang, Fengbin et al. 2019. “Structure of Microbial Nanowires Reveals Stacked Hemes That Transport Electrons over Micrometers.” Cell 177(2): 361–369.e10. https://linkinghub.elsevier.com/retrieve/pii/S0092867419302910.

Yalcin, Sibel Ebru et al. 2020. “Electric Field Stimulates Production of Highly Conductive Microbial OmcZ Nanowires.” Nature Chemical Biology 16(10): 1136–42. http://dx.doi.org/10.1038/s41589-020-0623-9.

Yoho, Rachel A., Sudeep C. Popat, and C??sar I. Torres. 2014. “Dynamic Potential-Dependent Electron Transport Pathway Shifts in Anode Biofilms of Geobacter Sulfurreducens.” ChemSusChem 7(12): 3413–19.

Zacharoff, Lori, Chi Ho Chan, and Daniel R. Bond. 2016. “Reduction of Low Potential Electron Acceptors Requires the CbcL Inner Membrane Cytochrome of Geobacter Sulfurreducens.” Bioelectrochemistry 107: 7–13. http://dx.doi.org/10.1016/j.bioelechem.2015.08.003.

Zhu, Xiuping, Matthew D. Yates, and Bruce E. Logan. 2012. “Set Potential Regulation Reveals Additional Oxidation Peaks of Geobacter Sulfurreducens Anodic Biofilms.” Electrochemistry Communications 22(1): 116–19. http://dx.doi.org/10.1016/j.elecom.2012.06.013.

