## Supplemental Information 1 for "Cytochrome expression shifts in *Geobacter sulfurreducens* to maximize energy conservation in response to changes in redox conditions"

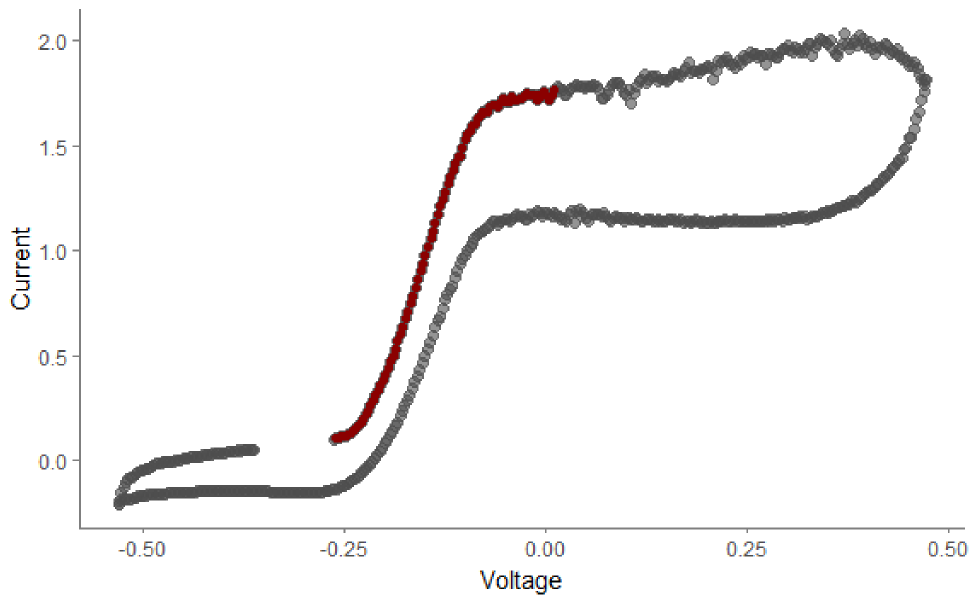


Figure S1: An example of a cyclic voltammogram with the model fitting region in red.

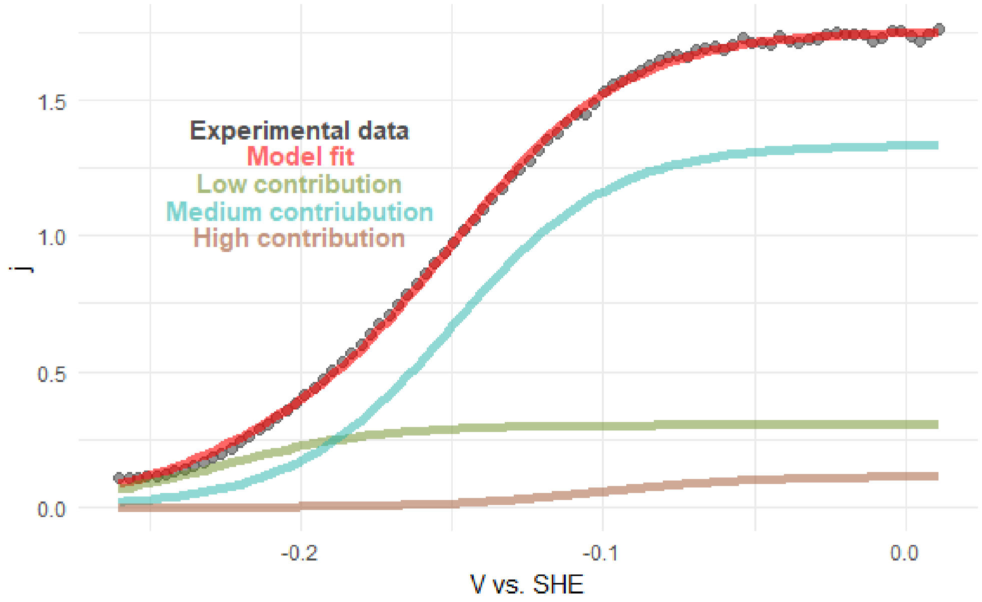


Figure S2: Graphical representation of the cyclic voltammogram with the model fit.

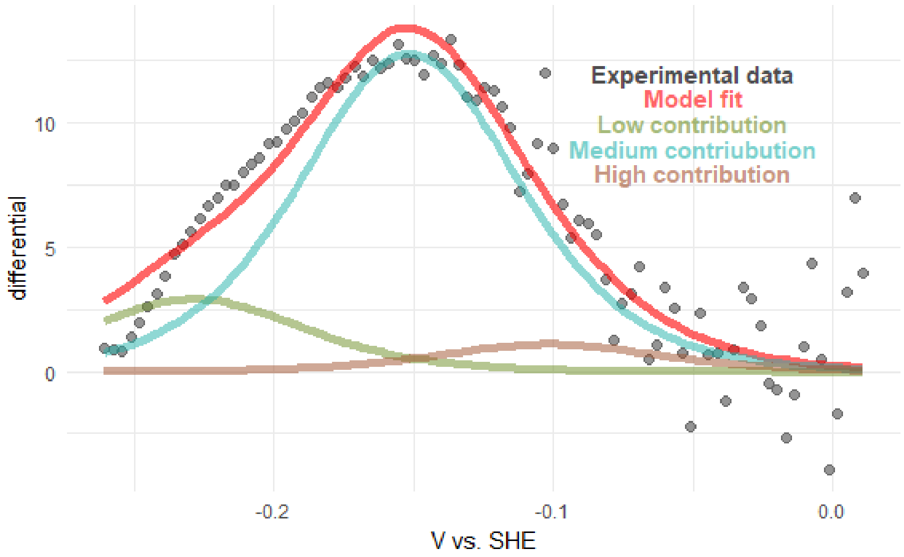


Figure S3: Derivation of the model fit and the differentiation of the cyclic voltammogram.
